# Arteriolar GMP signaling revealed by FRET-based real-time measurement in vivo

**DOI:** 10.64898/2026.05.26.728053

**Authors:** Lukas Thiele, Kjestine Schmidt, Martin Thunemann, Andreas Friebe, Susanne Feil, Robert Feil, Cor de Wit

## Abstract

cGMP evokes arteriolar vasorelaxation and is generated in smooth muscle by the soluble guanylyl cyclase (sGC) stimulated by endothelial NO. Phosphodiesterases (PDEs) degrade cGMP and may contribute in diameter regulation. We examined arteriolar cGMP levels in real-time *in vivo* upon NO and acetylcholine (ACh) and effects of PDE inhibition.

Changes in cGMP were measured by FRET using the indicator protein cGi500. Arterioles in the cremaster muscle of anesthetized mice expressing cGi500 were exposed for imaging. They were stimulated with various NO donors (DEA/NO, SNP, SNAP) and ACh with or without PDE inhibition (non-specific: IBMX; PDE-5 specific: sildenafil). Furthermore, dilations induced by ACh and SNP were studied in sGC-deficient mice.

DEA/NO and SNAP led to a fast, concentration-dependent rise of fluorescence ratio of cGi500 (by 4.0% at 10µM), that declined on removal slowly. These ratio changes indicated cGMP increases as verified by inhibition of the sGC using ODQ. PDE inhibition increased cGMP levels slowly despite NO synthase inhibition. However, it did not amplify NO-induced ratio increases, but the decline after drug removal was decelerated. Interestingly, ACh did not modify cGMP levels and dilations were not impaired in sGC-deficient mice.

FRET-based imaging allows reliable real-time assessment of arteriolar cGMP levels *in vivo*. PDE activity does not limit the amplitude, but the duration of cGMP signaling after stimulation. Furthermore, we conclude that ACh does not release endothelial NO in murine arterioles. Preliminary experiments demonstrate that simultaneous measurement of cGMP and diameter is feasible using mice with expression of cGi500 in smooth muscle cells.

## Introduction

The nucleotide cyclic guanosine monophosphate (cGMP) plays an important role in the regulation of arteriolar tone in the microcirculation. In vascular smooth muscle cells (VSMC), it activates the cGMP-dependent kinase type 1 (cGKI, also known as PKG1).^1^ This enzyme indirectly reduces the intracellular calcium concentration, leading to reduced activity of the myosin light chain kinase (MLCK). Additionally, it is reported that myosin light chain phosphatase (MLCP) activity is enhanced following cGKI activation.^2,3^ Both pathways result in dephosphorylation of the myosin light chain (MLC), thereby promoting relaxation of VSMC and thus vasodilation.^4^

Vascular cGMP can be generated by two enzymes, firstly, the particulate guanylyl cyclase (pGC), which is located in the plasma membrane and activated by natriuretic peptides such as ANP or CNP,^5^ and, secondly, the soluble guanylyl cyclase (sGC , also known as NO-GC), which is located in the cytosol of VSMCs and stimulated by nitric oxide (NO). The main source of vascular NO is the endothelium. It is produced by the endothelial NO synthase (eNOS) in response to multiple stimuli. The gold standard to elicit eNOS activation is the application of acetylcholine (ACh) to initiate endothelium-dependent vasodilation. Physiologically more important is the continuous mechanical stimulation of the endothelium by the flowing blood exerting a force onto the endothelial cell named shear stress. The resulting eNOS activation is responsible for the well-known endothelium-dependent flow-induced vasodilation.^6,7^

Phosphodiesterases (PDEs) degrade cyclic nucleotides by hydrolysis. In different tissues, 11 PDE families with distinct substrate specificities for cAMP and/or cGMP have been identified. In vascular tissue, the PDE-1, -3, -4 and -5 have been identified and, so far, PDE-4 and -5 demonstrated to be of functional importance.^8^ While PDE-1 and PDE-3 degrade both, cAMP and cGMP, PDE-4 is specific for cAMP, and PDE-5 preferentially degrades cGMP.^9,10^ Since cGMP inhibits PDE-3, elevation of cGMP concomitantly enhances cAMP levels leading to a synergistic vasodilatory effect of cAMP/cGMP elevating compounds.^11^ Reportedly, also PDE-9 may contribute to cGMP degradation in vascular smooth muscle cells, at least in culture.^12^

In the past, changes in the intracellular cGMP concentration were measured using different end-point methods that provide a snapshot and thus momentary information, e.g. cGMP levels acutely after stimulation. A well-established procedure is the radioimmunoassay (RIA) using antibodies directed against cyclic nucleotides on tissue preparations after NO donor application, thereby demonstrating increasing cGMP concentrations upon stimulation.^13,14^ Likewise, antibodies against cGMP used on fixed cells combined with immunohistochemistry allowed the localisation of cGMP increases in these experiments.^15^ However, both methods are unsuitable to continuously follow dynamic changes in the cGMP concentration over time in living cells and tissues in real time. Other investigators aimed for temporal resolution of cGMP changes in cells using a sophisticated method in which they employed a modified patch clamp method using cyclic nucleotide gated channels as a readout for cGMP levels.^16^

The development of fluorescent sensor proteins that respond differently to light after a conformational change has revolutionized the measurement of intracellular signaling molecules. Close distance between the two fluorescent domains enables radiation-free energy transfer (Förster Resonance Energy Transfer, FRET) and a conformational change with separation of the two domains abrogates this transfer. By tagging the cGMP-binding domain of cGKI with two fluorescent proteins, which are separated from each other upon cGMP binding, such a cyclic GMP indicator protein (cGi500) was generated^17^ which allows real-time cGMP measurements. Transgenic cGMP sensor mice expressing cGi500 can be used to monitor cGMP signaling in isolated primary cells or tissues or in *in vivo* models.^18^ Indeed, the use of cGMP biosensors has led to several conceptual advances, such as the discovery of local, intercellular and mechanosensitive cGMP signals.^19^

In this study, we aimed to demonstrate and quantify NO- and ACh-induced cGMP changes in the microcirculation of cGi500 sensor mice *in vivo*. To this end, we continuously monitored changes in light emission by cGi500 in the arteriolar wall of anesthetized mice. The role of distinct PDEs in cGMP degradation in this tissue was addressed by pharmacological inhibition of PDEs. Furthermore, we addressed the contribution of eNOS-independent mechanisms on basal cGMP levels upon PDE inhibition or sGC inhibition in the presence of a NOS inhibitor. Specifically in the microcirculation, the importance of NO as mediator of ACh-induced endothelium-dependent dilation is controversial.

Therefore, we also assessed the efficacy of ACh to induce cGMP increases in the vascular wall and the requirement of cGMP production for ACh-induced dilations.

## Material and methods

### Transgenic animals

Animal care and experiments were in accordance with the German Animal Welfare Act and approved by local authorities (Ministerium für Landwirtschaft, Umwelt und ländliche Räume des Landes Schleswig-Holstein). Male transgenic mice that expressed a cGMP indicator (cGi500) globally as previously described^18^ were examined at an age between 4 and 12 months. The cGi500 protein binds cGMP with an EC_50_ of about 470 nmol/L^20^ and allows relative cGMP measurement based on fluorescence resonance energy transfer (FRET). It consists of the cGMP binding domain of cGKI that is flanked by cyan fluorescent protein (CFP) and yellow fluorescent protein (YFP). Upon cGMP binding, FRET from CFP to YFP decreases resulting in a shift of the emitted light from 535 nm to 480 nm upon excitation at 425 nm.^21^ Arteriolar dilations were studied in mice that were globally deficient for the beta1 subunit of the sGC^22^ and respective wildtype littermates. In a single experiment, a conditional cGMP sensor mouse that expressed cGi500 in smooth muscle cells was studied. This was achieved by crossing a Nestin-Cre mouse line, expressing Cre in smooth muscle cells and/or their progenitors,^23,24^ to a Cre-inducible cGi500 mouse line.^18,25^

### Preparation of the microcirculation for intravital microscopy

Mice were anesthetized with intraperitoneal injection of fentanyl (0.06 mg/kg), midazolam (6 mg/kg), and dexmedetomidin (0.3 mg/kg) in a volume of 13 mL/kg. After insertion of a small tube into the jugular vein, the anesthetic drugs were continuously infused at a rate of 5 to 7 mL/kg/h (for 5 mL/kg/h: 0.02, 2.0, 0.10 mg/kg/h of fentanyl, midazolam, and dexmedetomidin). A small tube was inserted into the trachea to secure the airway and to ventilate the animal during the experiment with a tidal volume of 200 µl at a rate of 150 /min (MicroVent Mouse Ventilator, Hugo Sachs Elektronik, Hugstetten, Germany). The cremaster muscle was prepared as described^26^ and superfused with a tempered (34° C) buffer containing (in mmol/L): 118.4 NaCl, 20 NaHCO_3_, 3.8 KCl, 2.5 CaCl_2_, 1.2 KH_2_PO_4_, 1.2 MgSO_4_. The buffered (pH 7.4) saline solution was gassed with 5% CO_2_ and 95% N_2_ resulting in a pO_2_ of about 30 mmHg on the cremaster muscle due to contamination with ambient air. The muscle was superfused at a rate of 8 mL/min by hydrostatic pressure. Vasoactive drugs were added during the experiment to the superfusion buffer using a roller-pump (Minipuls 3, Gilson) at a ratio of 1:100 (or 1:50 for lipophilic drugs) through a side branch before the superfusion reached the cremaster muscle. Thereby and due to the high flow rate of the buffer, it was possible to add and remove the drug under investigation quickly at specific time points.

### Intravital FRET microscopy

A small number of arterioles was exposed in each animal for FRET microscopy by carefully removing the surrounding skeletal muscle for a length of about 2 mm using a surgical microscope (Wild MS-C, Wild-Heerbrugg AG, Switzerland). After surgery, the mouse was placed onto the microscope (Axioskop FS, Carl Zeiss AG, Göttingen, Germany) and the cremaster observed through a 40-fold objective (Achroplan 40x/0.75 water immersion lens, Carl Zeiss AG). The microscope was equipped with a commercially available system enabling FRET microscopy (Visitron Systems GmbH, Germany). The light source was either a polychromator system set to 420 nm (Visichrome, Munich, Germany) (data presented in figures 1, 2 and 5A) or a light-emitting diode (LED) with a light spectrum exhibiting a maximum at 425 nm (pE-100, CoolLED Limited) (for all other data). The latter was combined with a 425/26 nm filter (AHF, Tübingen, Germany) in the excitation light path to narrow the spectrum of the excitation light further. Excitation light was projected onto the specimen using a 450 nm dichroic mirror. Emitted light was projected simultaneously onto two separate cameras (Photometrics Evolve A16J103001) using a beam splitter (TwinCam) equipped with 505 nm dichroic mirror and 470/24 or 535/30 nm emission filters for CFP and YFP fluorescence detection, respectively. The hardware was controlled by software, which was also used to collect and store images (VisiView, Visitron Systems GmbH, Germany) on a PC. Images were sampled at a frequency of 1 Hz.

**Figure 1:**
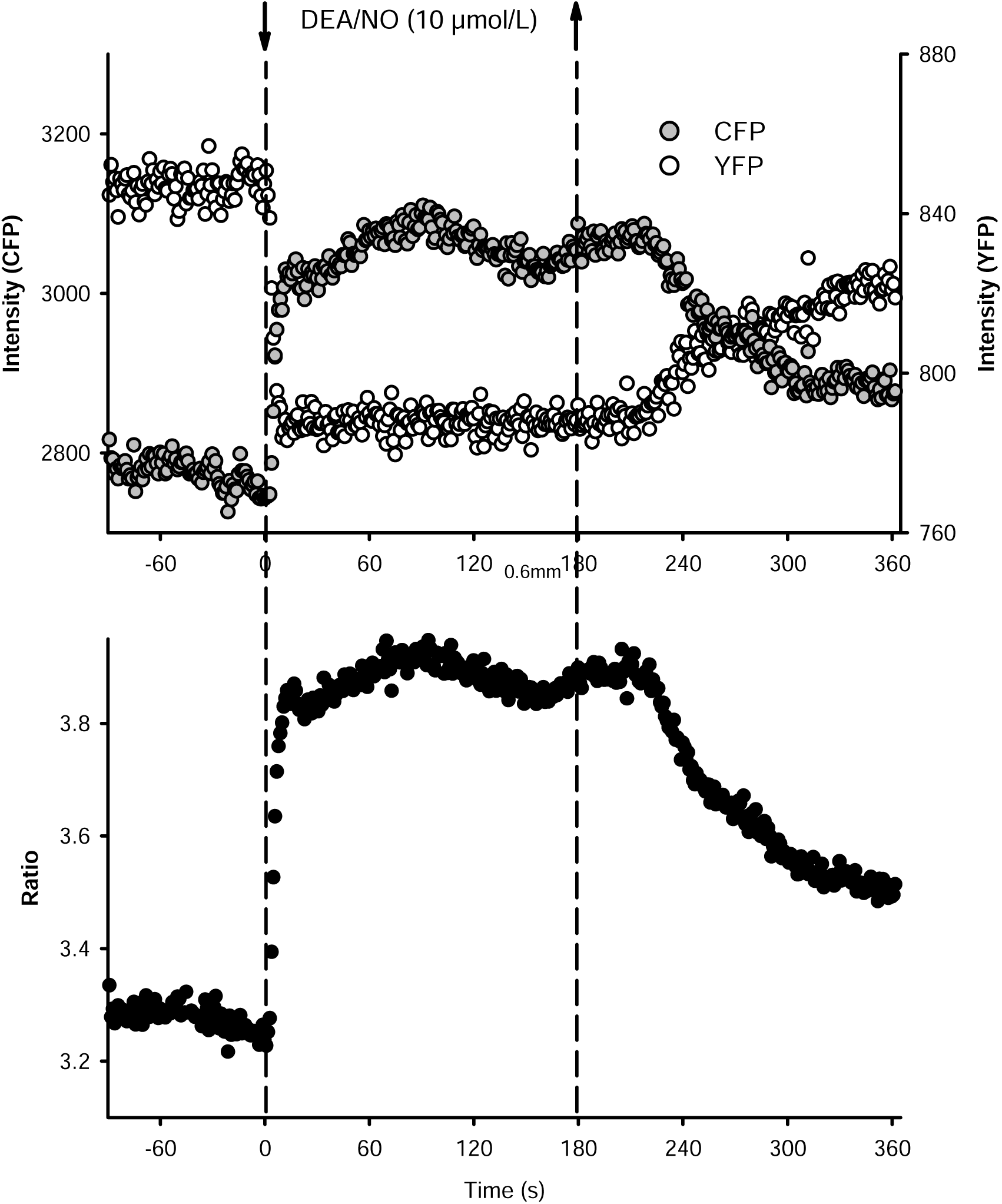
Fluorescence changes in an arteriole upon application of an NO-donor. Fluorescence changes in the vessel wall (upper panel) and the calculated ratio (lower panel) are plotted over time. The NO-donor DEA/NO (10 µmol/L) was applied at time 0 and removed after 180 s as indicated by the dashed lines with arrows. The fluorescence of the yellow fluorescent protein (YFP, open symbols) decreased immediately and, importantly, the fluorescence of the cyan fluorescent protein (CFP, grey symbols) increased. These fluorescence changes are indicative of a reduced energy transfer from CFP to YFP due to the conformational change of the cGi500 protein caused by enhanced cGMP- binding. Consequently, the ratio of the CFP and YFP fluorescence increases (lower panel).

**Figure 2:**
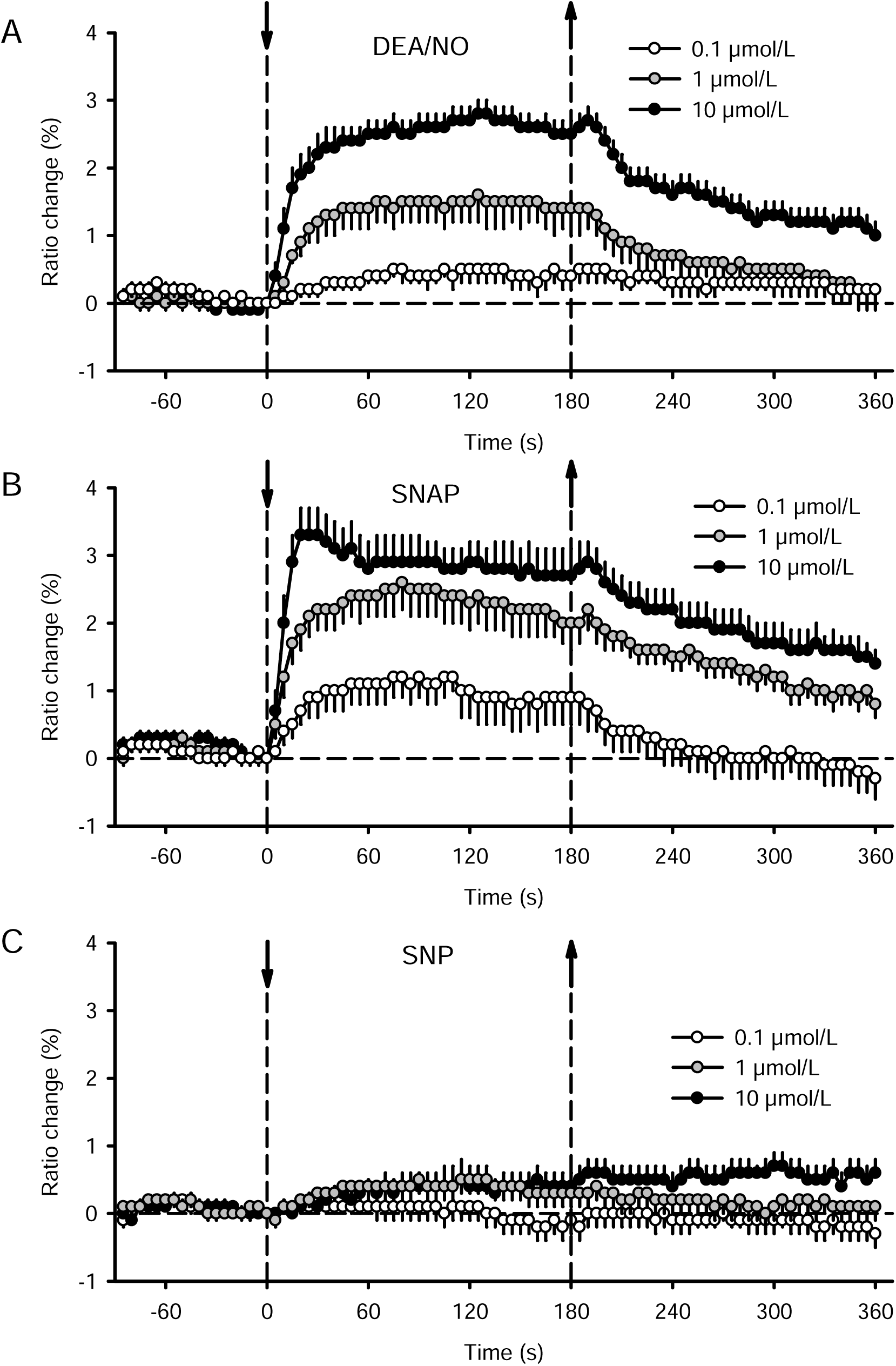
NO-donors induced concentration-dependent ratio increases. Superfusion of the NO-donors diethylamine-NONOate (DEA/NO, A), S-Nitroso-N-acetyl-DL- penicillamine (SNAP, B), and sodium-nitroprusside (SNP, C) induced concentration-dependent increases of the CFP/YFP ratio in the arteriolar wall which is indicative of enhanced cGMP levels. DEA/NO and SNAP induced comparable temporal responses although SNAP tended to act already at lower concentrations and the increases lasted longer after removal of the drug. SNP was much less efficient. NO-donors were applied for 180 s, starting at time point 0, as indicated by the dashed lines with arrows. Ratio is given as a percentage of the value measured immediately before drug application, n=15 to 17 arterioles in 7 mice, for statistical comparison of the mean values during application s. text.

The cGi500 mouse expressing the sensor in smooth muscle cells (cGi500^fl/fl^; Nestin-Cre) was studied using a spinning disk microscope as described.^25^ It comprised an upright microscope (Axio Examiner Z1, Zeiss), a Yokogawa CSU-X1 spinning disk confocal scanner, a DualView beam splitter with appropriate filters, and EM-CCD cameras (QuantEM 512, Photometrics) to record images of CFP and YFP emission simultaneously. A pE-2 LED system (CoolLED) was used for epifluorescence illumination. The system was controlled by VisiView software (Visitron Systems). Vascular diameter was measured from images collected by the software during ratio measurement.

### Experimental procedure for FRET/cGMP measurements

In each mouse, one to four arterioles were examined after equilibration of the preparation for 30 min. Only vessels with preserved blood flow were selected for the experimental procedure which was verified by inspection using the experimental microscope. Initially, the effect of different vasoactive drugs and concentrations were examined. The drugs were added to the superfusion fluid for 180 s while fluorescence was measured before and during drug application. Fluorescence measurements continued for at least another 180 s after removal of the drug. We examined different NO donors (diethylammonium-NONOate [DEA/NO], sodium-nitroprusside [SNP], S-nitroso-N-acetylpenicillamine [SNAP]), adenosine (a known stimulator of cAMP synthesis^27,28^), and the endothelium-dependent dilator acetylcholine (ACh). The NO-donor concentrations ranged from 0.1 to 10 µmol/L. In a subgroup of experiments, the different classes of vasoactive drugs were compared and, for this, a single, higher concentration was applied (10 µmol/L). After stimulation, the arteriole was allowed to recover for 3 to 10 minutes before the next concentration or drug was studied in the same vessel. Upon completion of all planned drugs at their respective concentrations in the arteriole, the next vessel was examined similarly. Thus, paired data were generated for different drugs and concentrations in a non-cumulative manner. For these experimental groups, the polychromator system was used as light source whereas in all other following groups the LED was used.

In further experimental series, the sGC as source of cGMP synthesis was blocked by 1H-[1,2,4]- oxadiazolo[4,3-a]-quinoxaline-1-one (ODQ, 30 µmol/L) or the degradation of cGMP was blocked by inhibition of PDEs. In different subsets of experiments two distinct PDE inhibitors were continuously applied, the non-specific inhibitor 3-isobutyl-1-methylxanthine (IBMX, 100 µmol/L) or the specific PDE5 inhibitor sildenafil (SIL, 1 µmol/L). In these experiments, the effects of NO donor application were examined at 0.01, 0.1, and 1 µmol/L during control condition in one to three arterioles and this was repeated in the same arterioles in the continuous presence of the respective inhibitor generating paired data. In a single group treated with IBMX, treatment order was reversed, i.e. initially, IBMX was continuously applied followed by its washout and examination of the responses without PDE inhibition. IBMX was dissolved as a stock solution in DMSO, therefore the solvent DMSO was applied and present at the same concentration (0.2%) during the respective control periods. These experimental series were performed in the continuous presence of the NOS inhibitor Nitro-*L*-arginine (L-NA, 30 µmol/L) to minimize any modulating effects of endogenous NO production. The animals were euthanized at the end of the experiments by an overdose of pentobarbital (2.5 g/kg) applied intravenously.

### Experimental procedure to study dilations

Intravital microscopy of the microcirculation to study dilations was described before.^29^ In brief, up to 13 arterioles were observed in each mouse through a 20-fold objective with a video camera-equipped microscope (Eclipse E600, Nikon, Germany). Microscopic images were displayed on a monitor at 700- fold magnification and recorded on videotape for later measurement of luminal diameter after digitization. The resolution after digitization was about 0.8 µm. Arteriolar diameters were measured before and during superfusion of different vasodilators (ACh, sodium nitroprusside, 8-Br-cGMP) at final concentrations from 0.01 to 10 µmol/L. After each concentration or substance, arterioles were allowed to return to their resting diameter for at least 4 min to assess non-cumulative concentration-response curves. Thus, non-stimulated resting diameters of all arterioles were measured manifold in each animal and the resting diameter of each arteriole is the mean of these multiple measurements. At the end of the experiment, the maximal diameter of the arterioles was determined by combined superfusion of ACh, adenosine and sodium nitroprusside (each 30 µmol/L). Animals were euthanized at the end of the experiment by intravenous application of pentobarbital.

### Data analysis

Fluorescence levels were determined from stored images using appropriate software (VisiView, Visitron Systems GmbH, Germany). Regions of interest were selected that represented the vascular walls. For these regions brightness was determined and values stored on PC for analysis using statistical software (STATA, Stata Corporation, Texas, USA). Ratio values were calculated by dividing brightness of the CFP channel by values from the YFP channel. These ratio values were used to calculate respective changes (% of control value) upon application of a drug.

Arteriolar diameter changes were normalized to the respective maximal possible response:

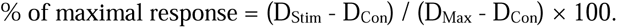

In this equation, D_Stim_ is the diameter after stimulation, D_Con_ is the control diameter before stimulation and D_Max_ is the respective maximal diameter for dilation and minimal for constriction observed during the experiment.

Statistical analysis was performed using commercially available software (STATA, College Station, Texas). Data are presented as a mean ± standard error of mean (mean ± S.E.M.). Paired data were sampled and paired comparison was performed before and during application of vasoactive drugs or for treatment groups without or with inhibitor (treatment vs control) during vasoactive drug application by paired t-test. Comparisons for diameter and diameter changes between different genotypes were performed using unpaired t-test. Differences were considered significant at an error probability of *P* < 0.05.

## Results

### Real-time measurement of cGMP in skeletal muscle arterioles in vivo

A total of 119 arterioles was studied in 43 mice. The skeletal muscle arterioles under study were second- (2A) and third-order arterioles (3A) which exhibit in this preparation typically maximal diameters in a range from 25 to 50µm. All mice expressed the cGMP-indicator protein cGi500 ubiquitously including the skeletal muscle in which the arterioles of the microcirculation are embedded. To avoid disturbance of the FRET measurements in arterioles by fluorescence from the skeletal muscle, arterioles were carefully separated from the skeletal muscle. This procedure led to loss of spontaneous vessel tone making it virtually impossible to measure changes of arteriolar diameter in response to vasodilator substances in experiments on mice with global cGMP sensor expression. As an example, fluorescence changes in the vessel wall and the calculated ratio change upon application of an NO donor (DEA/NO, 10 µmol/L) are displayed in figure 1. Upon superfusion of the NO donor, the fluorescence of the yellow fluorescent protein (YFP) decreased immediately and, importantly, the fluorescence of the cyan fluorescent protein (CFP) increased. These reciprocal fluorescence changes are indicative of a reduced energy transfer from CFP to YFP due to the conformational change of the cGi500 protein caused by enhanced cGMP-binding. Consequently, the ratio of the CFP and YFP fluorescence increases (Fig. 1) which is shown subsequently as a measure of intracellular cGMP levels.

### Concentration-dependent increases of cGMP in the arteriolar wall upon NO donors

Different NO donors applied at different concentrations onto the microcirculation induced reversible increases of the arteriolar cGMP concentration (Fig. 2). At the lowest concentration (0.1 µmol/L) DEA/NO increased the mean ratio measured during the application period of 180 s by 0.4±0.1% (p<0.05). This mean ratio increased further at 1 µmol/L to 1.3±0.3% (p<0.05 vs 0.1 µmol/L) and to 2.4±0.2% at 10 µmol/L (p<0.05 vs 1 µmol/L) showing a concentration-dependent increase of the cGMP level upon DEA/NO. The temporal behaviour is displayed in figure 2A. It shows a fast ratio increase within the first 15 s and a slower increase during a second phase for another 30 s before reaching a stable plateau that lasted until removal of the drug. Upon removal of DEA/NO, the ratio declined at a much slower rate as compared to the steep increase during the start of the stimulation. A different NO donor (SNAP) induced ratio increases comparable to DEA/NO, although the concentration-response was slightly shifted towards lower concentrations. SNAP induced larger increases already at the lowest concentration used (0.1 µmol/L: 0.9±0.3%, p<0.05) which was further enhanced to 2.2±0.3% at 1 µmol/L (p<0.05 vs 0.1 µmol/L). However, 10 µmol/L did not significantly increase the ratio further (2.7±0.3%, p=0.12 vs 1 µmol/L). The temporal pattern (Fig. 2B) was likewise comparable to DEA/NO with the exception that the highest concentration reached a plateau at an earlier time point. Together with the lack of a further ratio increase this indicates that maximal stimulation was achieved in this setting. In contrast, SNP was markedly less efficient to induce ratio increases (Fig. 2C). At 0.1 µmol/L no change was found (0.0±0.1%), whereas 1.0 µmol/L SNP induced a small increase in the mean ratio during its application (0.3±0.1%, p<0.05). The highest concentration of SNP (10 µmol/L) induced a comparable change (0.3±0.2%, p=0.06).

### NO donors stimulated sGC to achieve increases of cGMP

The sGC was blocked by superfusion of ODQ (30 µmol/L) in the presence of L-NA (30µmol/L) to minimize endogenous NO production. In each animal (n=5), only one arteriole could be monitored continuously during ODQ application. In these vessels, the ratio decreased within the first 180 s after application from 3.40±0.12 to 3.35±0.12 (by -1.5±0.1%, p<0.05). This observation was further substantiated in all vessels studied by comparing the ratios measured before the application of the different NO-donors in the absence or presence of QDQ: Ratio values before (3.49±0.07) were significantly higher than during ODQ application (3.41±0.05, p<0.05). As expected, DEA/NO (1 and 10 µmol/L) increased the ratio also in this experimental series and this increase was abrogated in the presence of ODQ (Fig. 3). Similarly, the ratio increase upon 1 µmol/L SNAP was abrogated and also strongly reduced upon 10 µmol/L SNAP (Fig. 3). However, a small, significant increase remained at this highest concentration.

**Figure 3:**
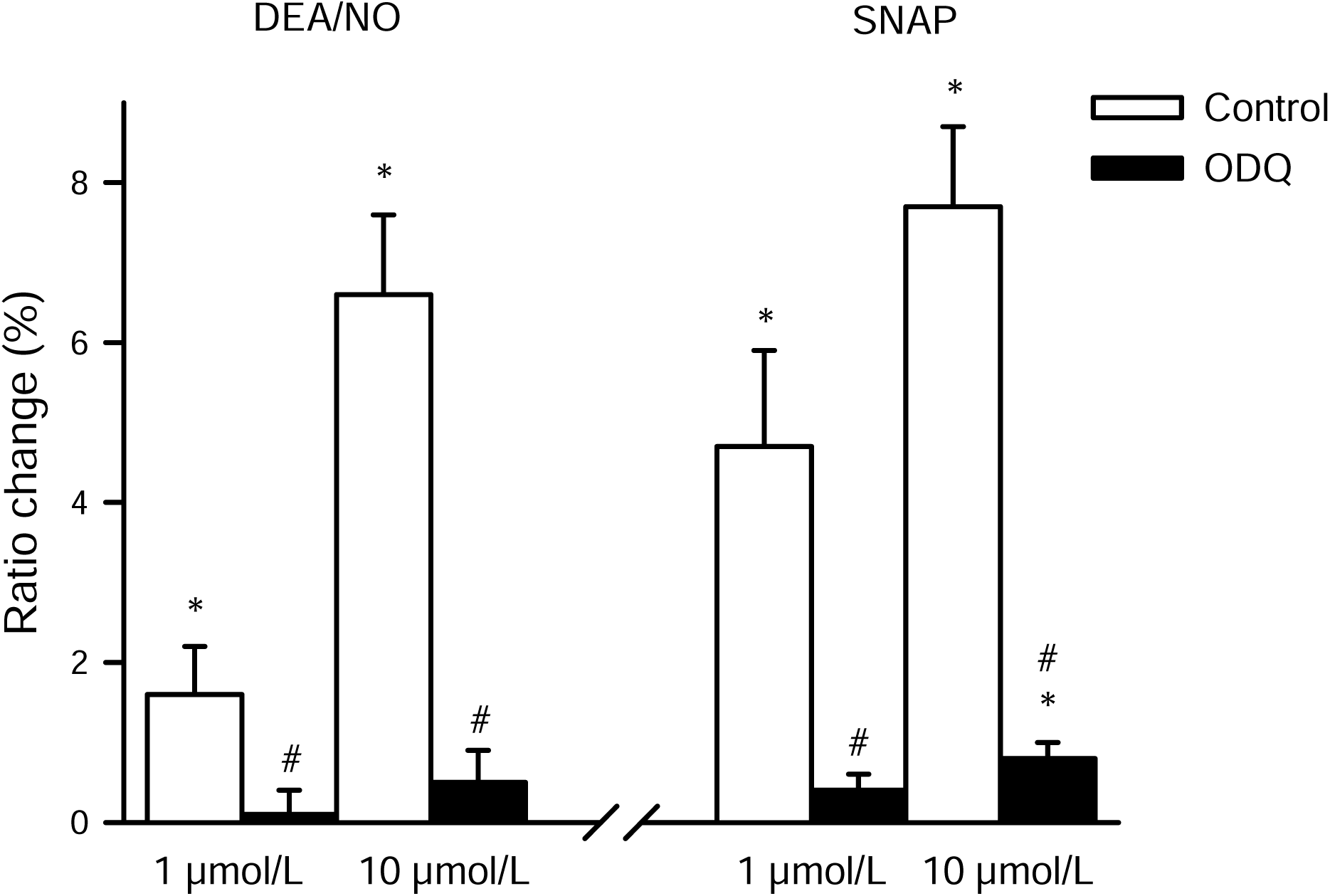
Blockade of the sGC abrogated cGMP increases in response to NO-donors. Mean value of the ratio changes measured during a 180 s period of application of DEA/NO and SNAP (each 1 and 10 µmol/L) are shown before and after addition of the sGC blocker ODQ (30 µmol/L). The ratio increases measured during control conditions in response to NO-donors (open bars) were completely abrogated in the presence of ODQ (black bars). Only for 10 µmol/L SNAP a small significant increase remained. n=15 arterioles in 7 mice, *: *P*<0.05 vs. value before respective NO-donor, #: *P*<0.05 vs. response during control conditions. All experiments were performed in the presence of L-NA (30 µmol/L).

*Inhibition of PDEs enhanced basal ratio levels in vessels but not NO-donor induced increases* Degradation of cGMP is achieved by the activity of phosphodiesterases (PDEs), which were inhibited non-specifically by IBMX (100 µmol/L). These experiments were done in the continuous presence of NO-synthase blockade (L-NA, 30 µmol/L). Upon application of the solvent of IBMX (DMSO, 0.2%) ratio values tended to decrease (from 3.39±0.07 to 3.35±0.08, n=15, p=0.06). In contrast, IBMX enhanced ratio values within the first 180 s after application from 3.37±0.12 to 3.43±0.12 (p<0.05, n=15), the increase over time is depicted in Figure 4A.

**Figure 4:**
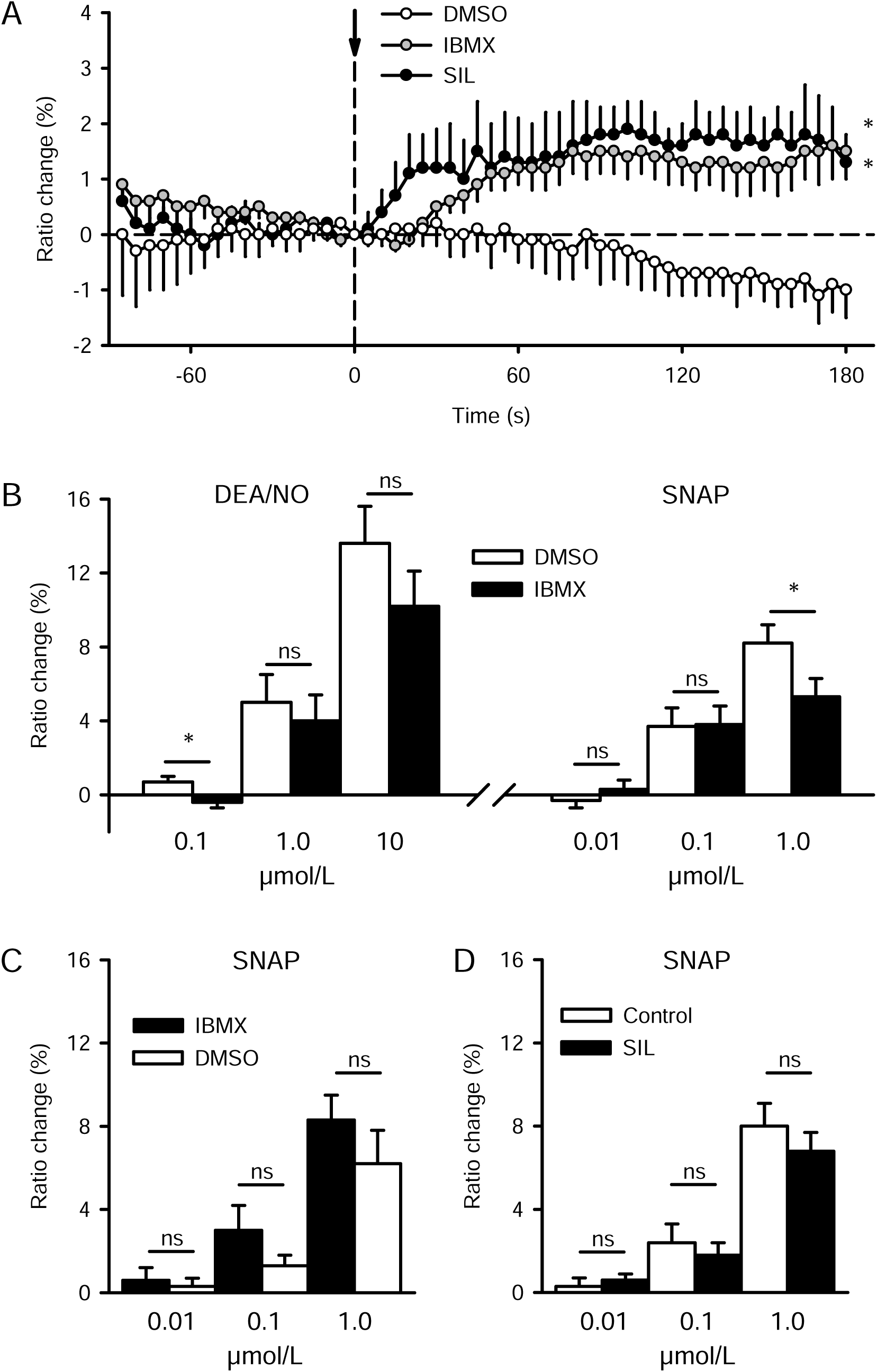
PDE inhibition enhanced basal cGMP levels in arterioles but not NO-donor-induced increases. A: Ratio changes are depicted over time before and after addition of the phosphodiesterase (PDE) inhibitors 3-isobutyl-1-methylxanthine (IBMX, 100 µmol/L) or sildenafil (SIL, 1 µmol/L) at timepoint 0 (indicated by the dashed line with arrow). As a control, the solvent of IBMX (DMSO, 0.2%) was applied. DMSO and IBMX: n=15 arterioles in 15 mice, SIL: n=6 arterioles in 6 mice, * indicates significant increase of the mean ratio value after application compared to initial value (*P*<0.05 ). B-D: Mean ratio changes during application of the respective NO-donor (applied for 180 s as shown in Fig. 2) in the absence or presence of PDE inhibitors. B: Control responses were examined firstly, followed by IBMX (DEA/NO or SNAP, n=15 arterioles in 5 mice each NO-donor). C: IBMX was applied first, followed by its washout (SNAP, n=15 arterioles in 5 mice). D: The effect of sildenafil was examined only for the NO-donor SNAP (n=18 arterioles in 6 mice). In none of these experiments (B to D) NO- donor-induced ratio changes were enhanced by PDE inhibition. ns indicates no significant difference, * significant difference (*P*<0.05 ) between NO-donor responses with or without PDE inhibition. All experiments were performed in the continuous presence of L-NA (30 µmol/L) to block NO-synthase.

**Figure 5:**
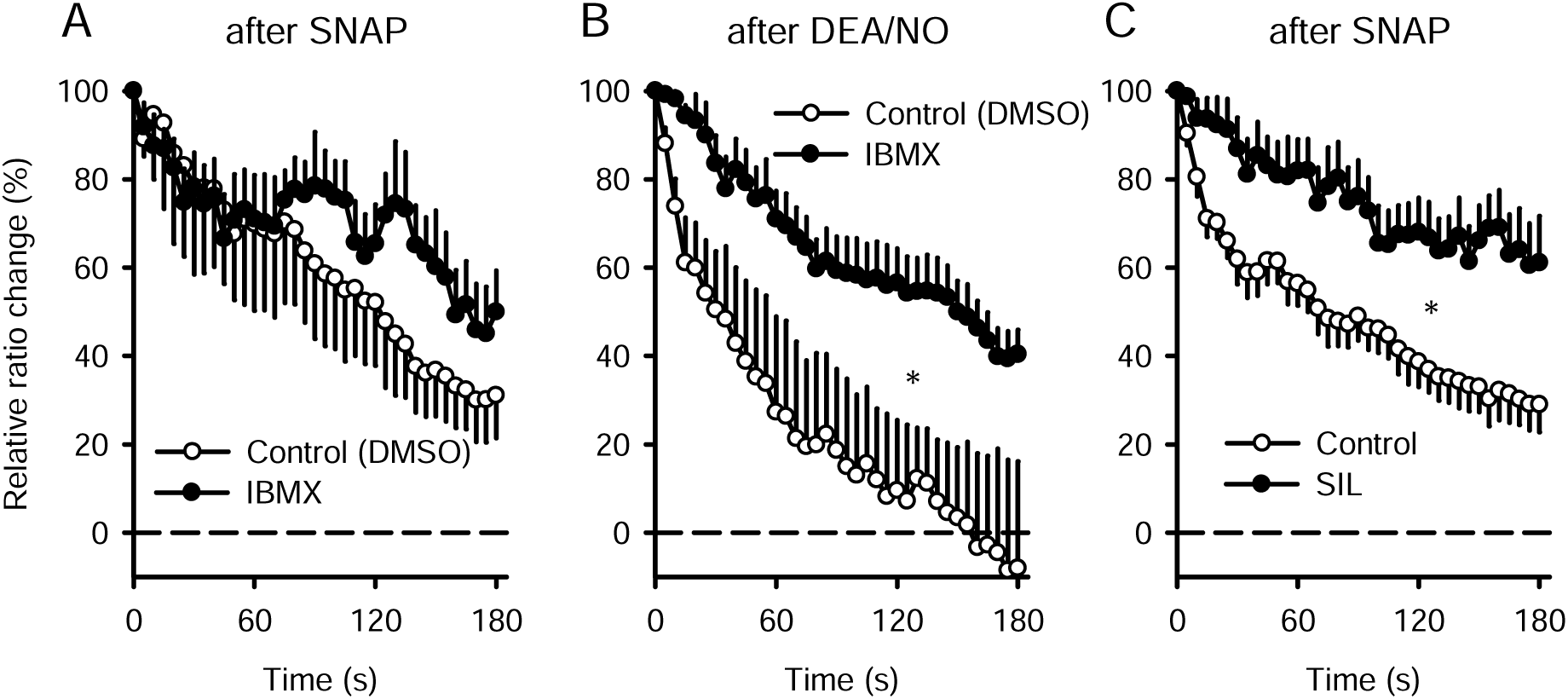
Ratio decrease after removal of NO-donors was delayed by PDE inhibition. The decline of the ratio values is depicted over time as a percentage of the increase observed before during NO-donor application. At timepoint 0 NO-donors (SNAP or DEA/NO, 1 µmol/L) were removed. Upon SNAP or DEA/NO removal ratio values declined towards initial values during control conditions (solvent DMSO 0.2% or water, open symbols). After DEA/NO ratio values declined faster and reached the initial level within 150s, while the ratio descent was slower after SNAP. IBMX did not modify the decline after SNAP (A), but significantly delayed the ratio reduction after DEA/NO (B). Sildenafil (SIL) also prolonged the ratio increase after SNAP (C). *: Significant difference between mean ratio value after NO-donor removal with or without PDE-inhibitor (*P*<0.05). Only vessels in which the ratio had increased (>1%) during NO-donor application were analysed (A: n=14, B: n=7, C: n=15 arterioles). All experiments were performed in the presence of L-NA (30 µmol/L).

The addition of the NO-donor DEA/NO increased ratio values in the presence of the solvent (DMSO) in a comparable manner as seen before, starting at 0.1 µmol/L with a subtle increase and up to 10 µmol/L with a stronger increase. In the presence of IBMX, DEA/NO-induced increases of the mean ratio values were, however, at neither concentration enhanced (Fig. 4B). Similarly, mean ratio changes induced by the application of SNAP were not enhanced in the presence of IBMX compared to the solvent alone (Fig. 4B). In this experimental series, we included a lower concentration of SNAP (0.01 µmol/L) that did not change the ratio during solvent application and it remained like this in the presence of IBMX. The unexpected inability of IBMX to enhance the ratio changes in response to NO-donors may be due to a desensitisation of the sGC or a negative feedback elicited by enhanced basal cGMP levels. Therefore, we performed an additional series of experiments in which we reversed the experimental protocol: At first, we examined SNAP responses in the presence of IBMX and thereafter re-examined SNAP responses after washout of IBMX in the presence of the solvent DMSO. Also, in this reversed series SNAP increased the mean ratio values during its application (at 0.1 and 1.0 µmol/L) in the presence of IBMX and during the subsequent control condition (DMSO). However, the increases were not significantly enhanced in the presence of IBMX (n=15, Fig. 4C).

In a different experimental series, the PDE5 was specifically inhibited by sildenafil (1 µmol/L). In each animal (n=6), only one arteriole could be monitored continuously during sildenafil application. Superfusion of sildenafil, in the presence of NO-synthase blockade (L-NA), enhanced ratio values within the first 180 s from 3.23±0.10 to 3.27±0.11 (p<0.05, n=6), the temporal behaviour of this increase (by 1.3±0.5%) is shown in Figure 4A. The observed increase of the ratio was sustained because it was also found in those vessels that were not continuously monitored. In these vessels, ratio values just before the application of different concentrations of SNAP during the ongoing experiment in the absence or presence of sildenafil were compared. Ratio values before (3.26±0.03) were significantly lower than in the presence of sildenafil (3.35±0.05, p<0.05, n=54 observations in 18 arterioles in 6 mice). While this indicates an efficient inhibition of PDE5, the ratio increases upon application of SNAP at different concentrations (including the very low concentration of 0.01 µmol/L) remained unaltered by sildenafil as compared to the respective control conditions (Fig. 4D).

### Inhibition of PDEs slowed down the ratio decrease after removal of NO-donors

Upon removal of the NO-donors, the ratio gradually declined and re-attained initial values (Fig. 2). We evaluated the effect of PDE-inhibition on the ratio decline upon removal of NO-donors by normalizing these values to the absolute increase observed during application. This was done for SNAP and DEA/NO (1 µmol/L) and only vessels were considered which responded with a ratio increase of more than 1% during NO-donor application. In such vessels, the mean relative ratio change upon SNAP washout was not significantly modulated by IBMX (59.2±14.3 vs. 69.8±6.6%, n=14, DMSO vs. IBMX, respectively). However, ratio decline after removal of DEA/NO was delayed in the presence of IBMX (DMSO: 23.2±16.9%, IBMX: 64.6±5.1%, n=7, p<0.05). In the presence of sildenafil, ratio decline was also retarded after removal of SNAP (Control: 48.3±4.7%, Sil: 74.8±6.0%, n=15, p<0.05). The decline over time of these groups is plotted in figure 5.

### Acetylcholine did not increase cGMP levels in arterioles

In addition to NO-donors, other vasodilators were studied in a separate experimental series. Expectedly, the NO-donors DEA/NO, SNAP, and SNP (each at 10 µmol/L) enhanced ratio values as seen before (mean ratio change: DEA/NO 3.7±0.8%, SNAP 3.3±0.6%, SNP 0.8±0.2%, all p<0.05, n=18). In marked contrast, acetylcholine (ACh, 10 µmol/L) did not significantly enhance ratio values in these same arterioles (1.0±0.6%, p=0.11, n=18). The mean ratio did also not change during superfusion of adenosine (10 µmol/L, 0.5±0.5%, p=0.37, n=18), which is known to dilate vessels via an increase of the cAMP concentration.^27,28^ All values before, during, and after agonist application are depicted in figure 6A.

**Figure 6:**
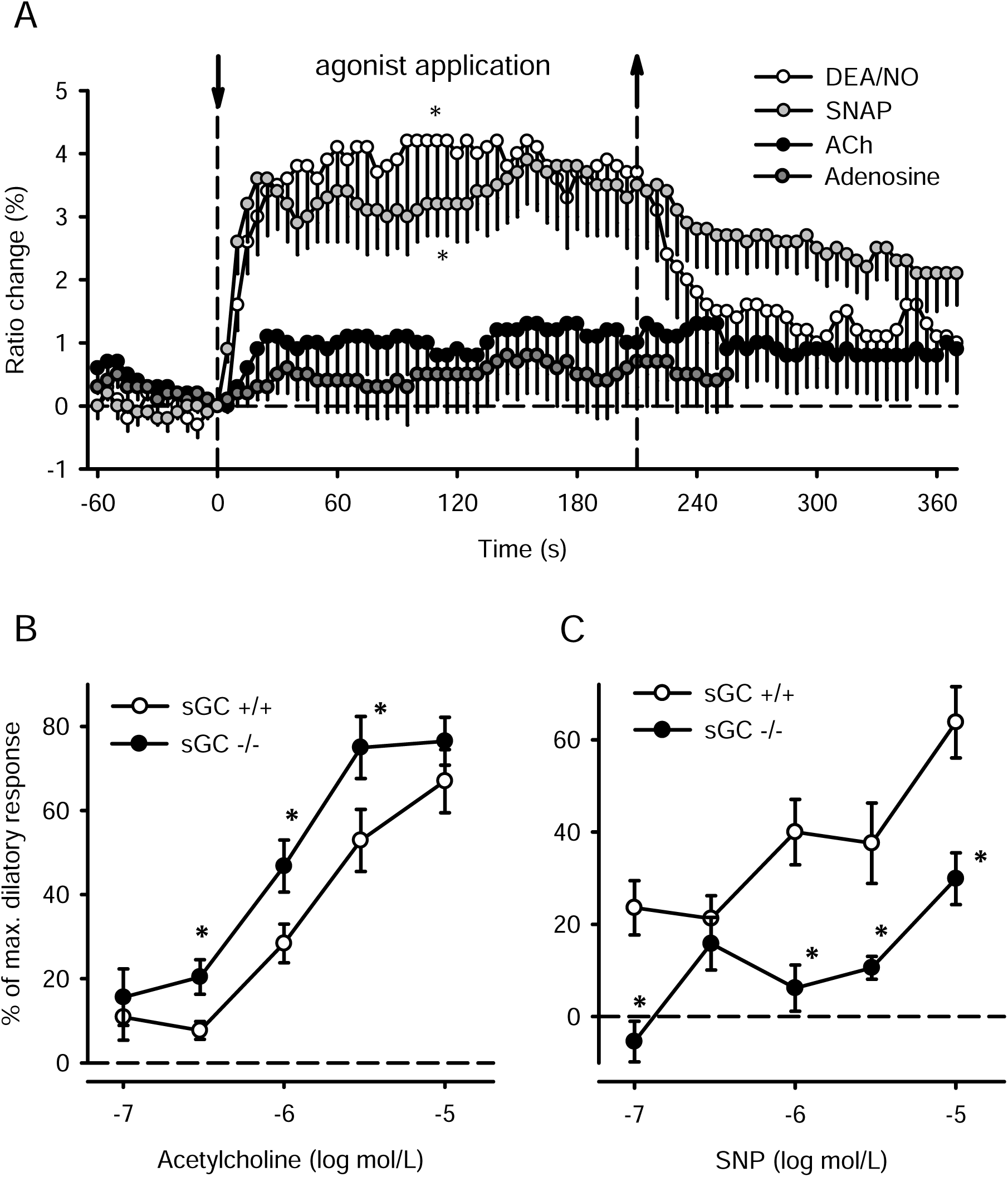
Acetylcholine (ACh) did not increase the CFP/YFP ratio and ACh-induced dilations are independent of sGC. A: The change of CFP/YFP ratio in response to superfusion of different vasoactive agonists (DEA/NO, SNAP, ACh, adenosine; each 10 µmol/L) is shown. Dashed lines with arrows indicate the start and the end of agonist application. DEA/NO and SNAP induced significant ratio increases, whereas ACh or adenosine did not modify mean ratio values significantly. n=18 arterioles in 6 mice, * indicates significant increase of mean ratio value during agonist application (*P*<0.05). B, C: Concentration-dependent arteriolar dilations in response to acetylcholine (B) or sodium-nitroprusside (C) in the microcirculation of wildtype (sGC^+/+^, open circles) and sGC-deficient mice (sGC^-/-^, black circles) are shown. SNP-induced dilations are abrogated, while ACh-induced dilations were not attenuated and even slightly enhanced in sGC-deficient mice. n=24 or 22 arterioles in 2 mice of each genotype, * indicates significant difference between genotypes (*P*<0.05).

### Acetylcholine-induced arteriolar dilations are independent of sGC

Dilations upon ACh were studied in wildtype and sGC-deficient mice (sGC^-/-^)^22^ in the microcirculation of the same skeletal muscle without arteriolar dissection. A total of 24 and 22 arterioles were studied in 2 mice of each genotype. The maximal diameter ranged from 13 to 59 µm and was not different between genotypes (n=24 with 33±2 and n=22 with 34±2 µm, wt and sGC^-/-^, respectively). However, resting diameters tended to be smaller and the quotient of resting and maximal diameter was lower in sGC^-/-^ arterioles (sGC^-/-^: 0.41±0.05, wt: 0.58±0.05, p<0.05) demonstrating a higher spontaneous tone in sGC^-/-^ arterioles compared to wildtype. ACh induced concentration-dependent dilations in both genotypes. The magnitude of the dilations was not attenuated in sGC^-/-^ arterioles, in fact, dilations were slightly stronger at intermediate concentrations (Fig. 6B). In marked contrast, concentration-dependent dilations induced by the NO-donor SNP were nearly abrogated in sGC^-/-^ arterioles (Fig. 6C). Dilations in response to 8- Br-cGMP (10 µmol/L) were, however, not reduced in sGC^-/-^ arterioles (wt: 31±8%, sGC^-/-^: 37±10%).

### Simultaneous measurements of cGMP and dilation is feasible

Finally, we examined the feasibility of simultaneous assessment of cGMP changes and dilations by measuring the CFP/YFP ratio as well as vascular diameter in the microcirculation. For this purpose, we examined a mouse which expressed the cGi500 sensor in smooth muscle cells, but not in skeletal muscle, using a Cre-inducible cGi500 mouse line and Cre expression controlled by the nestin promoter (cGi500^fl/fl^; Nestin-Cre). This approach avoids the otherwise required removal of the surrounding skeletal muscle to separate fluorescent signals originating from the arterioles. Thus, loss of spontaneous arteriolar tone and the resulting lack of dilation were avoided by the genetic targeting of the sensor. In this setting, the application of the NO donor DEA/NO increased CFP/YFP ratio and diameter almost simultaneously (Fig. 7). The removal of the NO donor induced a diameter decrease that preceded the return of the CFP/YFP ratio by 5 to 10 s (prominently observed at the lower DEA/NO concentration). Likewise diameter increased very shortly before ratio increases were detected by the cGi500 sensor. This suggests that the signaling pathway affecting the contractile machinery to induce diameter changes is more sensitive to changes of cGMP than the cGi500 sensor.

**Figure 7:**
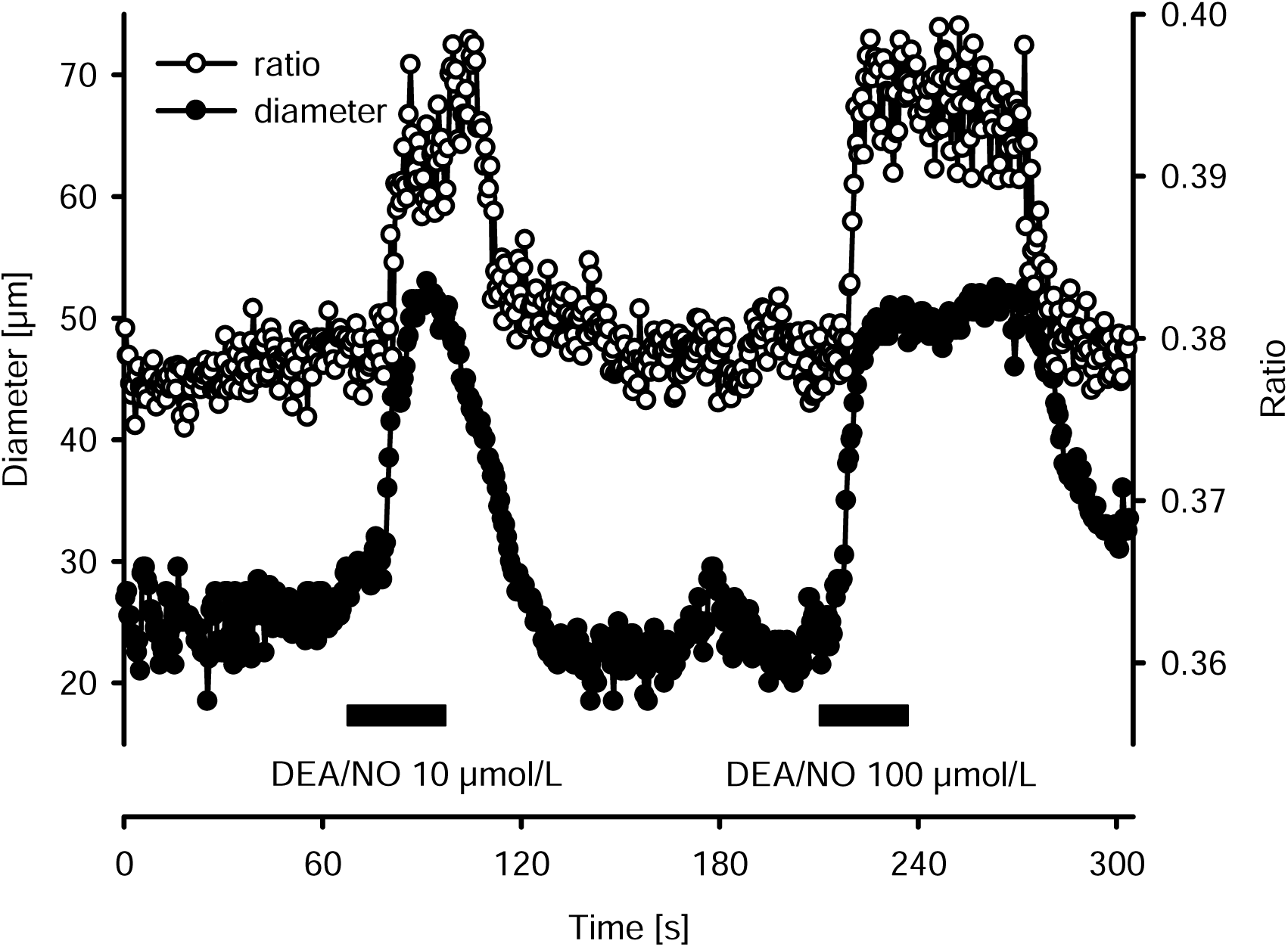
Simultaneous measurement of CFP/YFP ratio and diameter during NO-donor application. CFP/YFP ratio and diameter are shown as measured in an untouched arteriole that exhibited spontaneous tone in a mouse with cGi500 expression in VSMC, but not in skeletal muscle. The addition of the NO donor (DEA/NO, indicated by bar) induced near simultaneous increases of the CFP/YFP ratio and arteriolar diameter. Upon removal of the lower concentration of DEA/NO, diameter decreased before the ratio declined. In a similar fashion diameter increased slowly and very shortly before the ratio started to raise. This suggests that the signaling pathway affecting the contractile machinery is more sensitive to changes of cGMP than the cGi500 sensor.

## Discussion

This study demonstrates that FRET-based measurement of cGMP is a reliable method to follow changes in the cGMP concentration in the arteriolar wall in transgenic mice *in vivo* in real time. Exogenous NO induced rapid increases in the CFP/YFP ratio which were reversible upon removal of NO. These changes were indeed specific for changes in the cGMP concentration, as the ratio increases upon NO donors were abrogated after inhibition of sGC activity. Moreover, application of the cGMP-independent dilator adenosine did not alter the CFP/YFP ratio. While real-time ratio changes provide mechanistic insight, ratio measurements preclude the calculation of absolute cGMP concentrations, at least for intravital FRET measurements. Surprisingly, PDE inhibition did not further enhance ratio increases upon stimulation with NO, but it delayed cGMP decrement after NO removal.

The application of NO donors such as SNAP or DEA/NO led to prompt and concentration-dependent increases in the CFP/YFP ratio in the vascular wall. Among the used NO donors, SNAP was most potent and increased the ratio already at the low concentration of 0.1 µmol/L while for DEA/NO a concentration of 1 µmol/L was required to induce a comparable ratio increase. In addition, at higher concentrations, maximum levels were attained earlier when using SNAP. SNP, however, was less efficient. Cyclic GMP increases were detected within 5 seconds after NO donor application and final ratio levels were attained after 15 s, indicating a very rapid response. In a previous study using this sensor in a comparable approach for cGMP detection, the temporal resolution of cGMP increases in VSMC shortly after stimulation was not thoroughly analysed, but maximal responses apparently occurred only after minutes.^21^ This may be due to the fact that in the previous setup the stimulator hit the target only with a certain delay. However, cGMP increased within seconds in neuronal cells after NO donor application, as demonstrated by employing cyclic nucleotide gated channels as a readout for cGMP.^16^ Interestingly, flow-mediated vasodilation occurs within 60 s upon flow increase in conducting arteries.^6^ This is in line with the roughly 15 s lag for maximal cGMP increases upon NO application as observed in the present experiments considering other signaling mechanisms being additionally involved in the flow-mediated dilation.

After removal of NO donors, the ratio started to decline after 10 to 15 s and reached resting cGMP levels within 60 s at intermediate NO donor concentrations. However, at higher concentrations, the decline was slower and resting levels were not reattained within 3 min. In fact, the linear decline observed at high concentrations indicates that PDE degrades cGMP at maximum capacity in this setting, while at intermediate concentrations cGMP is initially degraded faster than at later time points.

To validate that changes in the CFP/YFP ratio reflect changes in the cGMP concentration, sGC was blocked by ODQ, a specific inhibitor of sGC.^30^ In the presence of ODQ, NO induced ratio changes were nearly abolished even at high NO donor concentrations. This provides evidence that ratio increases measured in the present experiment indeed mirror cGMP increases. Further, it demonstrates that NO activates sGC to increase cGMP levels. A conformational change of cGi500 may also be induced by cAMP binding. However, a substance known to enhance cAMP levels in VSMC (adenosine) did not enhance the CFP/YFP ratio even at high concentrations (10 µmol/L) providing further evidence for the specificity of cGi500 as previously demonstrated.^17^

Phosphodiesterases (PDE) such as PDE-5 degrade cGMP and thereby modulate the cGMP concentration in the arteriolar wall. Continuous endothelial NO release and resulting activation of sGC enhance cGMP levels, even more so during PDE inhibition. This may limit the detectable cGMP increase upon NO donor application. Thus, the NOS inhibitor L-NA was applied in all experiments studying PDE inhibition. Even during NOS blockade, non-specific PDE inhibition as well as specific PDE-5 inhibition evoked cGMP increases to a level comparable to those observed upon intermediate NO donor concentrations. However, this increase was slower and final cGMP concentrations were attained only after 60 s. The fact that PDE inhibition enhanced cGMP concentration even during NOS inhibition suggests that other mechanisms than local enzymatic NO production are responsible for basal cGMP production in the arteriolar smooth muscle *in vivo*. Either L-NA resistant NO production (insufficient NOS inhibition or NOS independent NO production), basal cGMP production by the sGC,^31^ or stimulation of the pGC by natriuretic peptides such as ANP^32^ may be responsible for those cGMP increases upon PDE inhibition. Moreover, NO may be liberated from S-nitrosothiols, nitrate, nitrite^33–35^ or released from erythrocytes entering the microcirculation with the blood stream. Erythrocytes express eNOS^36–38^ and their eNOS activity is likely maintained during L-NA application with the superfusion fluid. During local eNOS inhibition, NO derived from these sources enters the cremaster muscle arterioles, stimulates sGC and may enhance cGMP levels. This increase will be abrogated by local ODQ application, as was observed during NOS inhibition in the vascular wall.

One might anticipate that cGMP increases upon sGC stimulation are enhanced in the presence of PDE inhibition. Such a potentiation has been reported for isolated smooth muscle cells of different origin upon sGC stimulation using DEA/NO in the presence of IBMX.^18^ In the present experiments, the amplitude of NO induced cGMP increases, however, was not enhanced in the presence of IBMX or SIL. High NO donor concentrations may saturate cGi500, thereby preventing the detection of further cGMP increases. To circumvent a saturation of cGi500 in this experimental series, we applied low to intermediate NO donor concentrations and even concentrations that did not induce cGMP increases in the absence of PDE inhibition. Despite these precautions, we did not find an enhancement of the NO induced cGMP increases in the presence of IBMX for DEA/NO and SNAP or in the presence of sildenafil for SNAP. This demonstrates that degradation of cGMP by PDE is not a relevant mechanism limiting the amplitude of cGMP increases during continuous sGC stimulation. Upon NO removal, however, cGMP levels declined slowly reflecting cGMP degradation. This process was indeed decelerated in the presence of IBMX. More selective targeting of PDE-5 using sildenafil revealed a similar deceleration suggesting PDE-5 as the most relevant cGMP degrading enzyme in the arteriolar wall in mice.

In contrast to our findings, it was previously reported that PDE inhibition markedly enhanced cGMP increases upon NO donors or natriuretic peptides in smooth muscle tissues. Thuneman et al.^18^ compared cGMP increases in cultured primary VSMCs with or without PDE inhibition for a longer time starting with stimulation and lasting until complete remission of cGMP levels after removal of the stimulatory compound. The revealed potentiation of cGMP increases by PDE inhibition was mostly due to the persisting high cGMP concentration after drug removal, which reflects the retarded degradation of cGMP in the presence of PDE inhibitors. This indicates that also in isolated cells, as seen here in the intact animal, PDE inhibition is most efficacious upon cessation of sGC stimulation.

The FRET-based cGMP measurement system can also serve as an indicator of endogenous NO production as NO itself is difficult to quantify. It is commonly assumed, that ACh stimulates endothelial cells to release NO in large amounts in different vessels and species, thereby, inducing profound vasodilation. However, it is controversial whether NO also contributes to ACh-induced vasodilation in the microcirculation or other mechanisms are solely responsible.^39^ Specifically in the murine cremaster microcirculation, evidence was provided that vasodilation in response to ACh is elicited by activation of Ca^2+^-dependent K^+^-channels evoking endothelial hyperpolarization which is transferred to adjacent VSMC.^40,41^ Nevertheless, NO may concomitantly be released after ACh stimulation. In this case, cGMP increases upon ACh application should be detected easily using this setup as an indirect NO indicator. However, ACh at considerably high concentrations did not elicit cGMP increases while in the same arterioles NO donors robustly did. Importantly, ACh-dilations were not attenuated in mice deficient for sGC while dilations induced by the NO-donor SNP were strongly abrogated. Thus, we conclude that ACh does not initiate the release of relevant amounts of NO from endothelial cells in this preparation and the ACh-induced dilation is independent of cGMP production and activation of cGKI in VSMC. This conclusion is also consistent with our previous observation that arteriolar dilation induced by NO donors is abrogated in mice deficient for cGKI, the target of cGMP, while ACh-induced dilation remained fully intact in cGKI-deficient mice.^42^

A limitation of the present experimental approach is the difficulty of measuring cGMP and diameter simultaneously in global cGMP sensor mice. These mice express the cGi500 sensor in all tissues, including skeletal muscle cells. Consequently, isolation of the arterioles from the surrounding tissue was required for selective measurement of fluorescent signals originating from vascular tissue, which led to substantial loss of spontaneous vascular tone and lack of dilation. However, decent isolation of the arterioles facilitated the detection of changes in light emission in a specific region of the arteriolar wall providing more accurate measurement of ratio changes. The inconvenience of lack of tone can be circumvented by studying mice expressing this sensor in VSMCs but not in skeletal muscle as demonstrated in this study in preliminary experiments. This approach allows the correlation of changes in cGMP levels with vascular diameter and our initial data indicate that the signaling pathway affecting the contractile machinery may be more sensitive to changes of cGMP than the cGi500 sensor.

In conclusion, we demonstrated that FRET based cGMP imaging is a reliable method to study cGMP changes in the arteriolar wall in real-time *in vivo*. Intravital microscopy using fluorescent sensors provides mechanistic insights on intracellular signaling due to the high temporal resolution. Here, we found that PDE activity, specifically PDE-5, limits the duration of enhanced cGMP signaling after sGC stimulation. Furthermore, we conclude that ACh is not an activator of eNOS in the murine microcirculation. However, mechanical stimuli acting on endothelial cells such as shear stress likely induce continuous NO release. In the future, cGMP measurement using FRET may be established as a useful, reliable method for indirect NO detection and may reveal dynamic information for NO release in response to mechanical stimuli.

